# Bacterial community profiling highlights complex diversity and novel organisms in wildlife ticks

**DOI:** 10.1101/807131

**Authors:** Siobhon L. Egan, Siew-May Loh, Peter B. Banks, Amber Gillett, Liisa Ahlstrom, Una M. Ryan, Peter J. Irwin, Charlotte L. Oskam

**Affiliations:** Vector and Waterborne Pathogens Research Group, College of Science, Health, Engineering and Education, Murdoch University, Perth, Western Australia, Australia; School of Life and Environmental Sciences, The University of Sydney, Sydney, New South Wales, Australia; Australia Zoo Wildlife Hospital, Beerwah, Queensland, Australia; Bayer Australia Ltd, Animal Health, Pymble, New South Wales, Australia

**Keywords:** Microbiome, ticks, Ixodida, wildlife, marsupials, Anaplasmataceae

## Abstract

Ticks (Acari: Ixodida) transmit a greater variety of pathogens than any other blood-feeding group of arthropods. While numerous microbes have been identified inhabiting Australian Ixodidae, some of which are related to globally important tick-borne pathogens, little is known about the bacterial communities within ticks collected from Australian wildlife. In this study, 1,019 ticks were identified on 221 hosts spanning 27 wildlife species. Next-generation sequencing was used to amplify the V1-2 hypervariable region of the bacterial 16S rRNA gene from 238 ticks; *Amblyomma triguttatum* (n=6), *Bothriocroton auruginans* (n=11), *Bothriocroton concolor* (n=20), *Haemaphysalis bancrofti* (n=10), *Haemaphysalis bremneri* (n=4), *Haemaphysalis humerosa* (n=13)*, Haemaphysalis longicornis* (n=4), *Ixodes antechini* (n=29), *Ixodes australiensis* (n=26), *Ixodes fecialis* (n=13), *Ixodes holocyclus* (n=37), *Ixodes myrmecobii (*n*=1), Ixodes ornithorhynchi* (n=10), *Ixodes tasmani* (n=51) and *Ixodes trichosuri* (n=3). After bioinformatic analyses, over 14 million assigned bacterial sequences revealed the presence of recently described bacteria ‘*Candidatus* Borrelia tachyglossi’, ‘*Candidatus* Neoehrlichia australis’, ‘*Candidatus* Neoehrlichia arcana’ and ‘*Candidatus* Ehrlichia ornithorhynchi’. Furthermore, three novel Anaplasmataceae species were identified in the present study including; a *Neoehrlichia* sp. in *I. australiensis* and *I. fecialis* collected from quenda (*Isoodon fusciventer*) (Western Australia), an *Anaplasma* sp. from one *B. concolor* from echidna (*Tachyglossus aculeatus*) (New South Wales), and an *Ehrlichia* sp. from a single *I. fecialis* parasitising a quenda (WA). This study highlights the diversity of bacterial genera harboured within wildlife ticks, which may prove to be of medical and/or veterinary importance in the future.

## 1. Introduction

Current estimates suggest that approximately 17% of all infectious diseases of humans are vector-borne (Rinker et al., 2016) and global trends show that vector-borne diseases (VBDs) are rising at a rapid rate (Jones et al., 2008; Morens and Fauci, 2012). The complex interplay between pathogen, vector, host(s) and the environment make VBDs particularly challenging to understand. In addition, factors such as climate change (Ostfeld and Brunner, 2015), land use (Ferrell and Brinkerhoff, 2018), feral animal populations (Merrill et al., 2018) and the microclimate within a landscape (Dobson et al., 2011) can further influence the prevalence and distribution of VBDs.

Ticks (Acari: Ixodida) comprise a group of haematophagous (blood feeding) arthropods with over 900 species described globally (Guglielmone et al., 2014; Mans et al., 2019). Ticks are known to transmit various pathogens; however, they also harbour a range of endosymbiont and commensal species (Špitalská et al., 2018). The epidemiology of recognised tick-borne diseases (TBDs) in the northern hemisphere demonstrates that wildlife serve as sentinels and can be used to monitor the presence and distribution of tick-borne pathogens (TBPs). Importantly, research has shown that some wildlife species act as dilution hosts for certain TBPs, whereas others may act as amplification hosts (LoGiudice et al., 2003).

The complexity of TBDs means that studies are increasingly shifting away from isolated species-specific studies toward ecosystem-based, collaborative research (Estrada-Peña et al., 2013; Pfäffle et al., 2013). Metabarcoding provides an informative molecular tool to characterise the bacterial diversity in ticks. Worldwide, next-generation sequencing (NGS)-based analyses have been applied to a range of tick species that are important from medical and veterinary perspectives, including *Amblyomma americanum* (Ponnusamy et al., 2014), *Ixodes ricinus* (Bonnet et al., 2014), and *Rhipicephalus microplus* (Andreotti et al., 2011).

Metabarcoding studies of the bacterial microbiome of Australian ticks have been reported only recently, with the first bacterial profiling by NGS conducted on the human-biting tick *Ixodes holocyclus* (Gofton et al., 2015a). These authors identified a highly abundant endosymbiont ‘*Candidatus* Midichloria mitochondrii’ (CMm) and after blocking the amplification of this organism, a greater bacterial diversity was revealed, including a number of novel microbes (Gofton et al., 2015a,b). Critically, in contrast to many parts of the world where multiple TBPs have been elucidated within well-studied tick-host-environment ecologies, there is a relative dearth of such information available for Australia. With this in mind, the aims of our study were to survey the bacterial communities present in ticks collected from Australian wildlife and to investigate their genetic relatedness to ‘taxa of interest’, i.e. tick-associated pathogenic and endosymbiotic organisms (Parola and Raoult, 2001; Mediannikov and Fenollar, 2014; Sumrandee et al., 2016). ‘Taxa of interest’ in the present study were defined as genera within alphaproteobacteria, gammaproteobacteria and spirochaetes known to be transmitted by ticks in other parts of the world, specifically; *Anaplasma*, *Bartonella*, *Borrelia*, *Coxiella*, *Ehrlichia*, *Francisella*, *Midichloria*, *Neoehrlichia*, *Rickettsia* and *Rickettsiella*. (Ahantarig et al., 2013; Vayssier-Taussat et al., 2015; Bonnet et al., 2017; de la Fuente et al., 2017).

## 2. Materials and methods

### 2.1 Sample collection and identification

1,019 ticks were sourced opportunistically from wildlife in Australia by veterinarians, veterinary nurses, wildlife carers, researchers and via submissions from members of the public, and preserved in 70% ethanol before being shipped to Murdoch University, Western Australia (WA), for identification. Ticks were identified morphologically to life stage and species using keys and species’ descriptions (Roberts, 1970; Barker and Walker, 2014). Details of sample collection are available in Supplementary File S1.

### 2.2 DNA Extraction

A sub-sample of ticks (*n*=238) was chosen for DNA extraction and bacterial profiling. Ticks were selected to represent as many tick species, hosts, geographical regions and life stages as possible within the study. Prior to DNA extraction, individual ticks were surface-sterilised in 10% hypochlorite solution, rinsed in 70% ethanol and DNA-free PBS, and then air-dried. Total genomic DNA (gDNA) was extracted using the Qiagen DNeasy Blood and Tissue kit (Qiagen, Germany) following the manufacturer’s recommendations with the following modifications; ticks were placed in a 2 mL safe lock Eppendorf tube with a 5 mm steel bead, frozen in liquid nitrogen for 1 minute and homogenised by shaking at 40 Hz in a Tissue Lyser LT (Qiagen, Germany). The volume of elution buffer AE was adjusted to 200 μL for engorged females, 100 μL for unengorged adults and 40 μL for nymphs. A double elution was carried out to increase gDNA yield for unengorged adults and nymphs. Sterile and DNA-free equipment and tubes were used for each step. Extraction reagents blank (EXB) controls were performed alongside tick extractions to assess background bacterial communities.

### 2.3 16S amplification and library preparation for NGS sequencing

Amplicons targeting a 250-320 bp product of the V1-2 hypervariable region of the 16S rRNA (16S) gene were generated using the primer pair 27F-Y (Gofton et al., 2015b) and 338R (Turner et al., 1999). Previous research identified that paralysis tick, *I. holocyclus*, has a highly abundant bacterial species (‘*Ca*. M. mitochondrii’), which masks the diversity of bacteria in 16S metabarcoding studies (see Gofton et al. 2015a). Therefore, a blocking primer developed by Gofton et al. (2015a) was used to inhibit the amplification of ‘*Ca*. M. mitochondrii’ in *I. holocyclus* ticks. Depending on tick life stage and level of engorgement, 3-10 μM of the CMm blocking primer was added to the PCR reaction. Amplicon PCRs were conducted in 25 μL reactions each containing 1 X Buffer (KAPA Biosystems, USA), 1.5 mM MgCl_2_, 0.4 μM of each forward and reverse primer with MiSeq adapters, 0.4 mg/mL BSA (Fisher Biotech, Australia), 0.25 mM dNTPs (Fisher Biotech, Australia), 0.5 U KAPA Taq (KAPA Biosystems, USA), and 2.0 μL of undiluted genomic DNA. Samples underwent the following thermal cycling conditions; initial denaturation at 95°C for 5 mins, followed by 35 cycles of denaturation at 95°C for 30 s, annealing at 62°C (CMm blocking primer present) or 55°C (CMm blocking primer absent) for 30 s, and extension at 72°C for 45 s with a final extension at 72°C for 5 mins. Libraries were then prepared using the Nextera XT DNA library preparation kit in 25 μL reactions following manufacturer’s recommendations. Libraries were purified using Agencourt^®^ AMPure^®^ XP PCR purification beads (Beckman Coulter Life Sciences, USA) and pooled in equimolar amounts. Final libraries were then quantified using the Qubit^®^ 2.0 Fluorometer (Thermo Fisher, Australia). Libraries were sequenced on an Illumina MiSeq™ using v2 chemistry (2×250 paired end).

Extraction reagents blanks (n=12) and PCR no-template controls (NTC; n=7) were included in all stages of the workflow. All pre- and post-PCR procedures were performed in physically separate dedicated laboratories and sterile protocols were maintained through library preparation in order to minimise amplicon contamination.

### 2.4 Bioinformatics and statistical analysis

Raw fastq files were downloaded from the Illumina BaseSpace Sequence Hub for analysis in a custom pipeline using USEARCH (Edgar, 2010). Raw paired-end sequences were merged in USEARCH v10, a minimum of 50 nucleotide (nt) overlap and maximum number of mismatches increased to 15 nt due to long overlap of paired-end sequences. Only sequences with perfect primer sequences were retained, primer sequences and distal bases were trimmed using USEARCH v8.0. Sequences were quality filtered in USEARCH v10, allowing a <1% expected error rate and singletons were discarded (Edgar and Flyvbjerg, 2015). Sequences were then clustered into operational taxonomic units (OTUs) of 97% similarity using the UPARSE algorithm (Edgar, 2013) in USEARCH v10. Taxonomy was assigned in QIIME2 v2018.4 using the q2-feature-classifier (Bokulich et al., 2018) with reference to a trained Greengenes database (DeSantis et al., 2006) (release May 2013) using the primer pair 27F-Y/ 338R. Taxonomic assignments were confirmed using NCBI MegaBLAST (Morgulis et al., 2008) on a random subsample of OTUs and in the case of tick-associated microbes, GenBank accession numbers and percentage identity of top hits were recorded for ‘taxa of interest’. The profiles from EXB controls and NTCs were first assessed to ensure quality of sampling and absence of tick-associated bacteria. The following criteria were used to assess inclusion of sequences: all OTUs that appeared exclusively in controls were removed; OTUs that had a higher relative sequence abundance in controls compared to tick samples were removed; and OTUs were removed that appeared in over half of controls (i.e. at least eight) that had a taxonomic identity associated with environmental bacteria (e.g. members of the phyla Acidobacteria and Cyanobacteria). In addition, potential cross-contamination during library preparation or ‘cross-talk’ at the sequencing level was assessed by inspecting the presence of expected tick-associated bacteria (e.g. members of the obligate intracellular bacterial families Anaplasmataceae and Midichloriaceae) in controls. The profiles of EXB controls and NTCs were then removed bioinformatically from associated samples to eliminate background bacteria.

Data analysis and visualisation was carried out in RStudio (RStudio Team, 2015) using packages metacoder (Poisot et al., 2017), phyloseq (McMurdie and Holmes, 2013) and vegan (Oksanen et al., 2019). Alpha diversity of samples was measured using observed OTUs, Chao1 index, Shannon index and Simpson index. Removal of samples with a low sequencing depth (<1000 assigned OTUs after data filtering), did not significantly alter alpha diversity measurements among tick species (data not shown). Rarefaction curves for samples were calculated to assess sequencing depth based on observed number of OTUs. Principal coordinate analysis (PCoA) on weighted unifrac dissimilarity measurements was used to assess the differences in microbial composition between tick species. Investigation into ‘taxa of interest’ warranted a more rigorous assessment of sequence number than was required for diversity measures in order to avoid any potential cross-contamination and machine cross-talk (Dong et al., 2017; Wang et al., 2017), as such only samples with >100 sequences were considered positive. Taxonomic assignment to ‘taxa of interest’ was assessed based on NCBI MegaBLAST top hit of named organism. Where precent identity was ≤97%, the terminology of nearest named Genus-like was employed. Due to sample collection biases in the present study, prevalence data of these microbes were not considered statistically relevant.

Nucleotide sequences from ‘taxa of interest’ were aligned by MUSCLE (Edgar, 2004) using default parameters and aligned sequences were then imported into MEGA7 (Kumar et al., 2016) with the most appropriate nucleotide substitution model chosen based on the lowest Bayesian Information Criterion (BIC) score. Evolutionary histories were inferred using the Neighbour-Joining method based on the Tamura 3-parameter model (Tamura, 1992). Bootstrap analysis was conducted using 10,000 replicates to assess the reliability of inferred tree topologies.

## 3. Results

### 3.1 Tick-host associations

Ticks were collected from 221 wildlife hosts including 24 native and three introduced species (Fig. 1, Supplementary File S1). Hosts recorded in the present study included; agile antechinus (*Antechinus agilis*), brown antechinus (*Antechinus stuartii*), spotted-tail quoll (*Dasyurus maculatus*), water rat (*Hydromys chrysogaster*), quenda (*Isoodon fusciventer*), northern brown bandicoot (*Isoodon macrourus*), western grey kangaroo (*Macropus fuliginosus*), eastern grey kangaroo (*Macropus giganteus*), red-necked wallaby (*Notamacropus rufogriseus*), red kangaroo (*Osphranter rufus*), platypus (*Ornithorhynchus anatinus*), eastern barred bandicoot (*Perameles gunnii*), long-nosed bandicoot (*Perameles nasuta*), sugar glider (*Petaurus breviceps*), eastern ring-tailed possum (*Pseudocheirus peregrinus*), black fruit-bat (*Pteropus alecto*), fruit-bat sp. (*Pteropus* sp.), bush rat (*Rattus fuscipes*), black rat (*Rattus rattus*), Tasmanian devil (*Sarcophilus harrisii*), wild pig (*Sus scrofa*), short-beaked echidna (*Tachyglossus aculeatus*), rufous-bellied pademelon (*Thylogale billardierii*), short-eared brush-tailed possum (*Trichosurus caninus*), common brush-tailed possum (*Trichosurus vulpecula*), wombat (*Vombatus ursinus*), red fox (*Vulpes vulpes*), and swamp wallaby (*Wallabia bicolor*). Female ticks were the dominant life stage recorded (*n*=547) followed by nymphs (*n*=319) and males (*n*=153). Ticks were received from animals in the Northern Territory (NT) (*n*=16), Queensland (QLD) (*n*=249), New South Wales (NSW) (*n*=316), Tasmania (TAS) (*n*=136), Victoria (VIC) (*n*=44) and Western Australia (WA) (*n*=258). Together, *Bothriocroton concolor* (*n*=123), *Ixodes australiensis* (*n*=210), *I. holocyclus* (*n*=173) and *Ixodes tasmani* (*n*=184) accounted for over two-thirds of all ticks submitted.

**Fig. 1.**
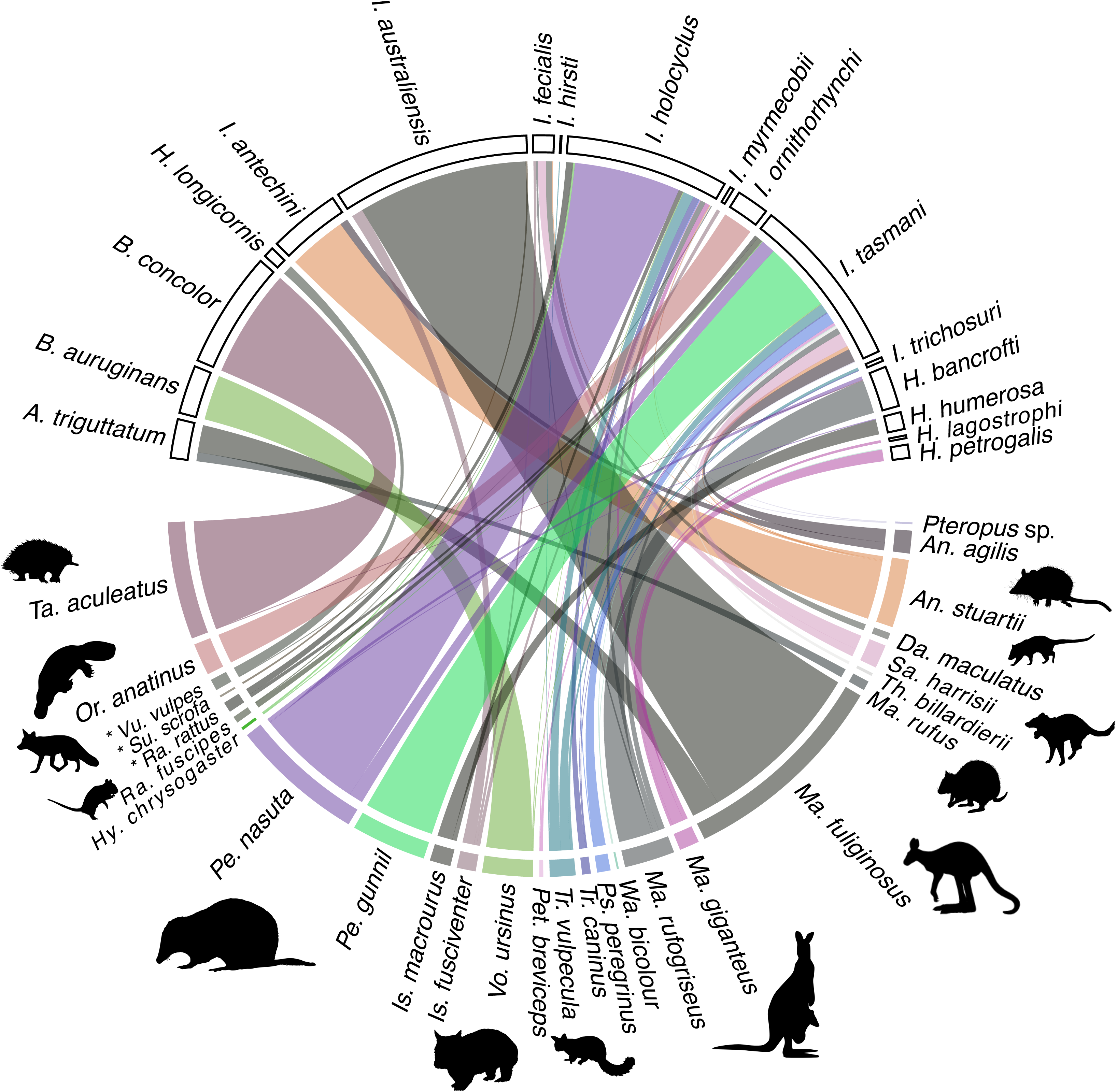
Chord diagram showing tick-wildlife associations recorded in the present study. The wildlife hosts recorded are represented on the lower half of the plot, and tick species across the top. Thickness of bar relative to number of ticks. Host records that could not be assigned to species level are not represented in this plot. A complete list of records is available in Supplementary File S1. Introduced wildlife species are denotes with an asterisk. Silhouette wildlife images sourced from phylopic.org.

### 3.2 16S rRNA bacterial profiling

A sub-sample of 238 ticks and 19 controls underwent 16S NGS profiling. A total of 23.9 million raw paired-end sequences were generated on the Illumina MiSeq. 17.9 million sequences were retained after merging, and subsequent quality filtering yielded 14.8 million sequences for clustering and taxonomic assignment. A total of 4,864 OTUs (average length 299 bases) were retained after background profiles were removed. After removal of background sequences, a total of 14,328,059 bacterial sequences were assigned to tick samples. Despite a high number of OTUs, only 1,535 OTUs had greater than 100 total sequences from tick samples. Tick samples had an average of 60,201 assigned sequences (see Supplementary File S2). *Amblyomma triguttatum* had the highest median alpha diversity as measured by the observed number of OTUs and the chao1 index, whereas *Ixodes antechini* had the highest median alpha diversity as measured by the Shannon and Simpson indexes (Fig. 2). Rarefaction analysis of the sequence depth shows that the observed number of OTUs plateaued at 50,000 sequences (Fig. 3). After the removal of sequences from controls, eight phyla were retained. An ordination plot of OTUs (Fig. 4) show that bacteria belonging to the Proteobacteria phylum were the most abundant and diverse taxa classified, followed by Firmicutes and Actinobacteria. Bacterial families identified in tick species, represented as relative number of sequences in Fig. 5, show 26 dominant taxa. While bacterial composition varied between tick species, sequences from the members of Coxiellaceae, Francisellaceae, and Rickettsiaceae families (Phylum: Proteobacteria) were the most abundant. Although sample sizes varied among tick species, beta diversity analysis incorporating abundance (sequence number) and taxonomic relatedness, showed evidence of different bacterial communities among tick species (Supplementary File S3).

**Fig. 2.**
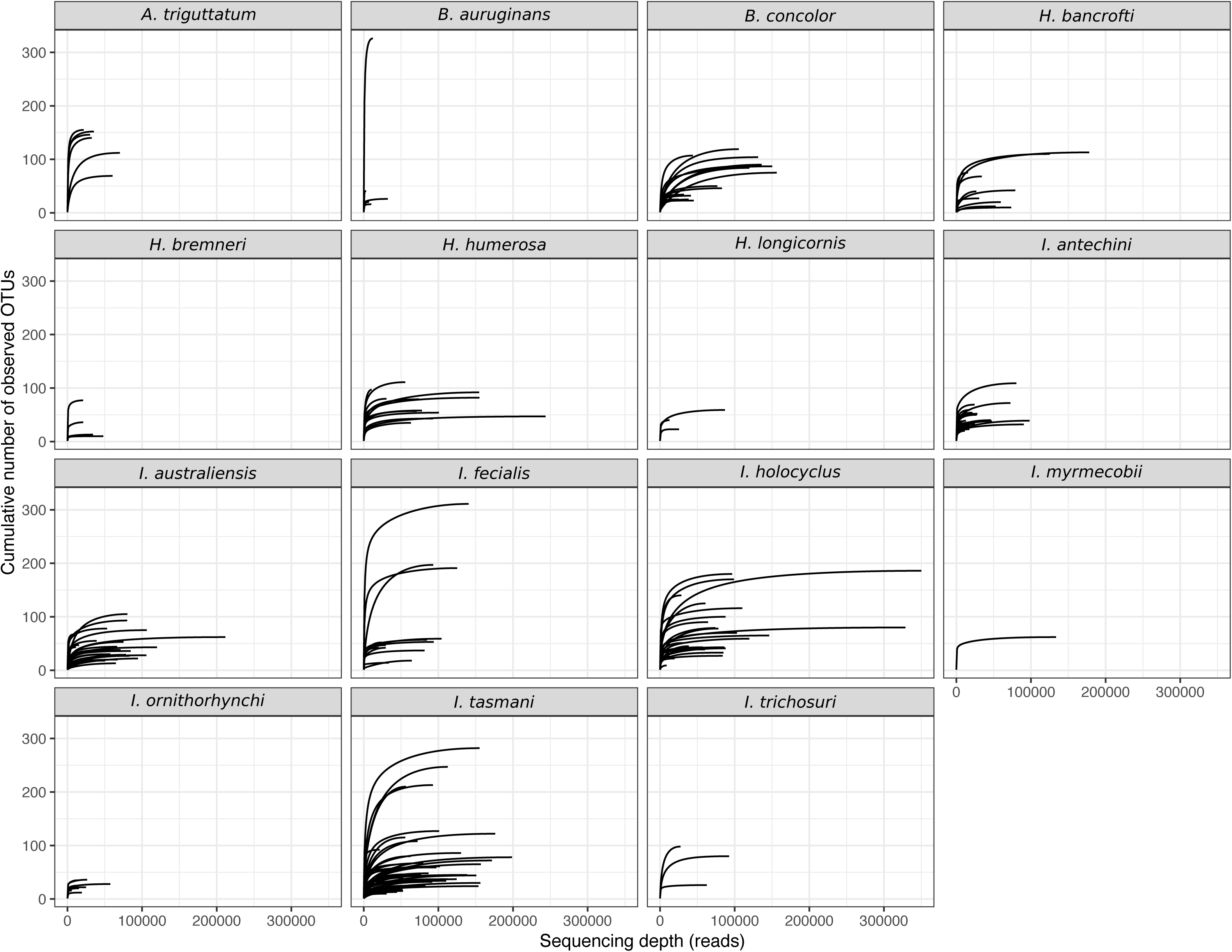
Alpha diversity of bacterial communities in ticks parasitising Australian wildlife measured as; observed number of operational taxonomic units (OTUs), Chao1 index, Shannon index and Simpson index. Tick species abbreviated to; *A. triguttatum* (*A. tri*), *B. aurugians* (*B. aur*), *B. concolor* (*B. con*), *H. bancrofti* (*H. ban*), *H. bremneri* (*H. bremneri*), *H. humerosa* (*H. hum*), *H. longicornis* (*H. lon*), *I. antechini* (*I. ant*), *I. australiensis* (*I. aus*), *I. fecialis* (*I. fec*), *I. holocyclus* (*I. hol*), *I. myrmecobii* (*I. myr*), *I. ornithorhynchi* (*I. orn*), *I. tasmani* (*I. tas*), and *I. trichosuri* (*I. tri*).

**Fig 3.**
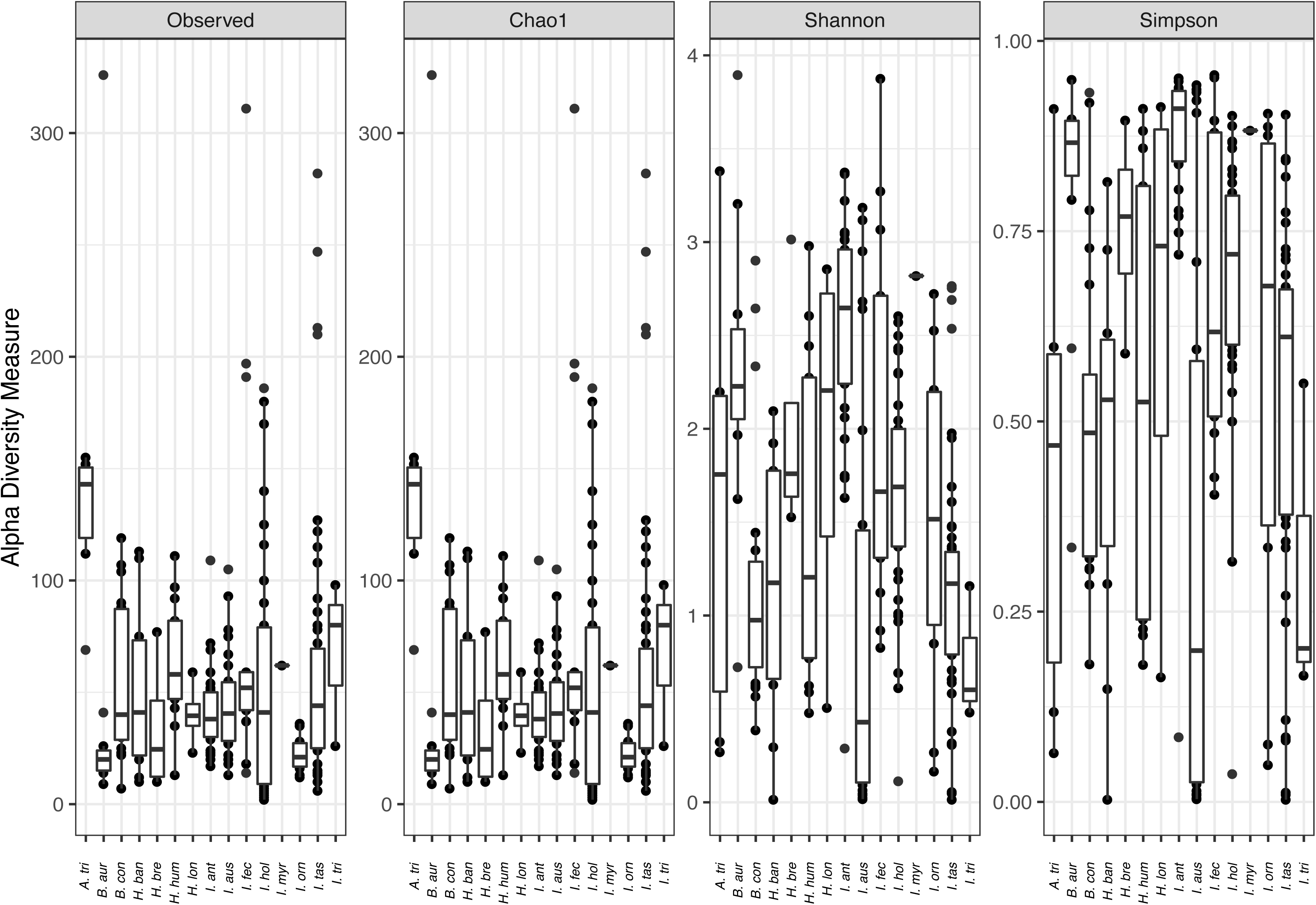
Rarefaction plot of 16S rRNA bacterial sequences clustered into 97% operational taxonomic units (OTUs) from ticks parasitising Australian wildlife.

**Fig 4.**
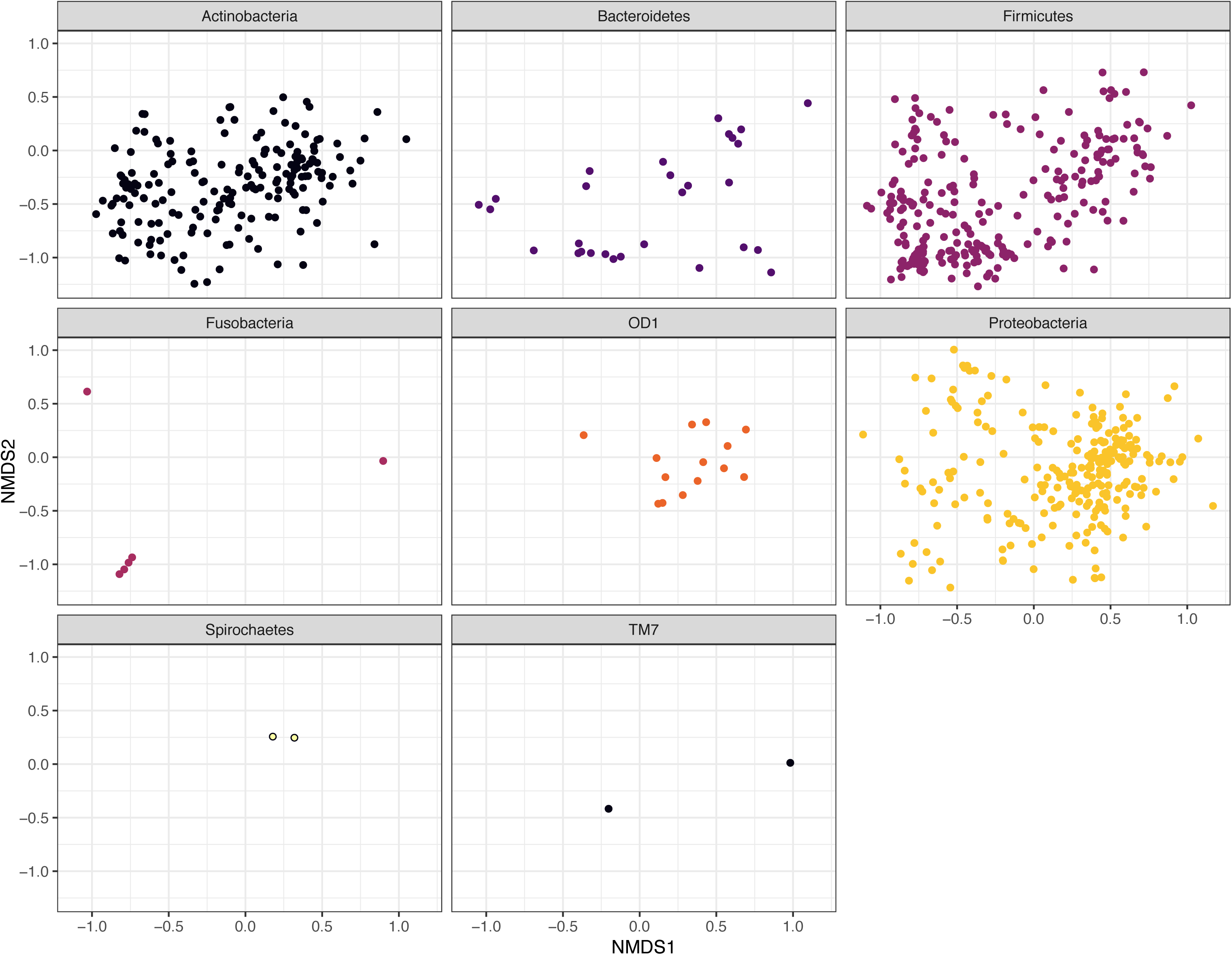
Non-metric multidimensional scaling (NMDS) plot of operational taxonomic units (OTUs) based on Bray-Curtis dissimilarity matrix. Taxa filtered to only display OTUs that were present at least twice in >10% of tick samples.

**Fig 5.**
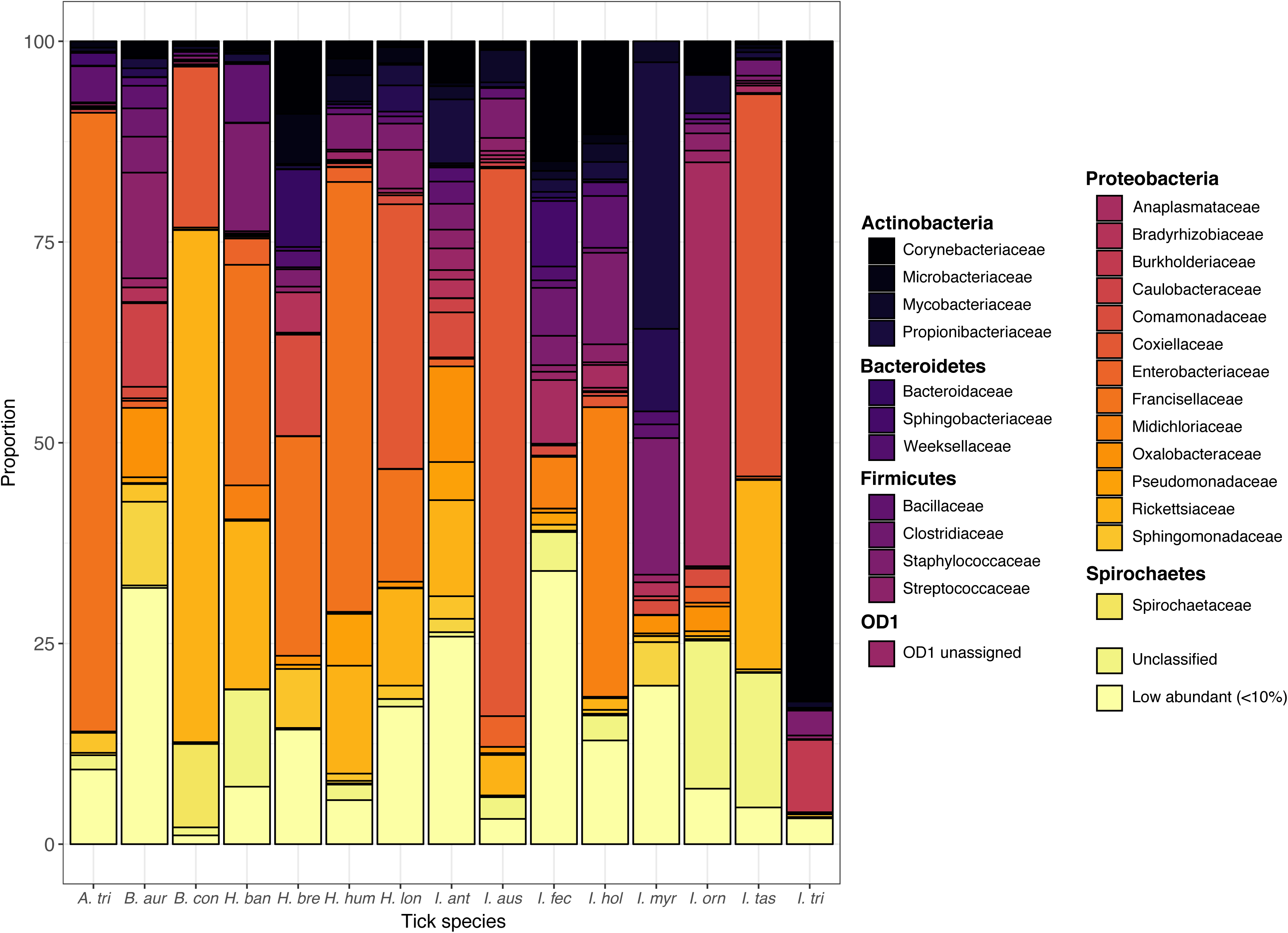
Stacked barplot of bacterial composition (shown at family level) from Australian ticks represented as percentage of assigned sequences. Taxa at the family level that represented <10% of the relative sequences within each tick species were grouped as “Low abundant”. Sequences not able to be accurately assigned to family taxa are displayed as “Unclassified”.

### 3.3 Presence of ‘taxa of interest’

In total, 37 OTUs were identified as ‘taxa of interest’ in the present study. Their taxonomic identity and abundance (number of sequences) in each sample is available in Supplementary File S4 (see Supplementary File S5 for fasta file of sequences). Phylogenetic analysis of the ‘taxa of interest’ is represented in Fig. 6, showing tick species and Australian states and territory, where each OTU was identified.

**Fig 6.**
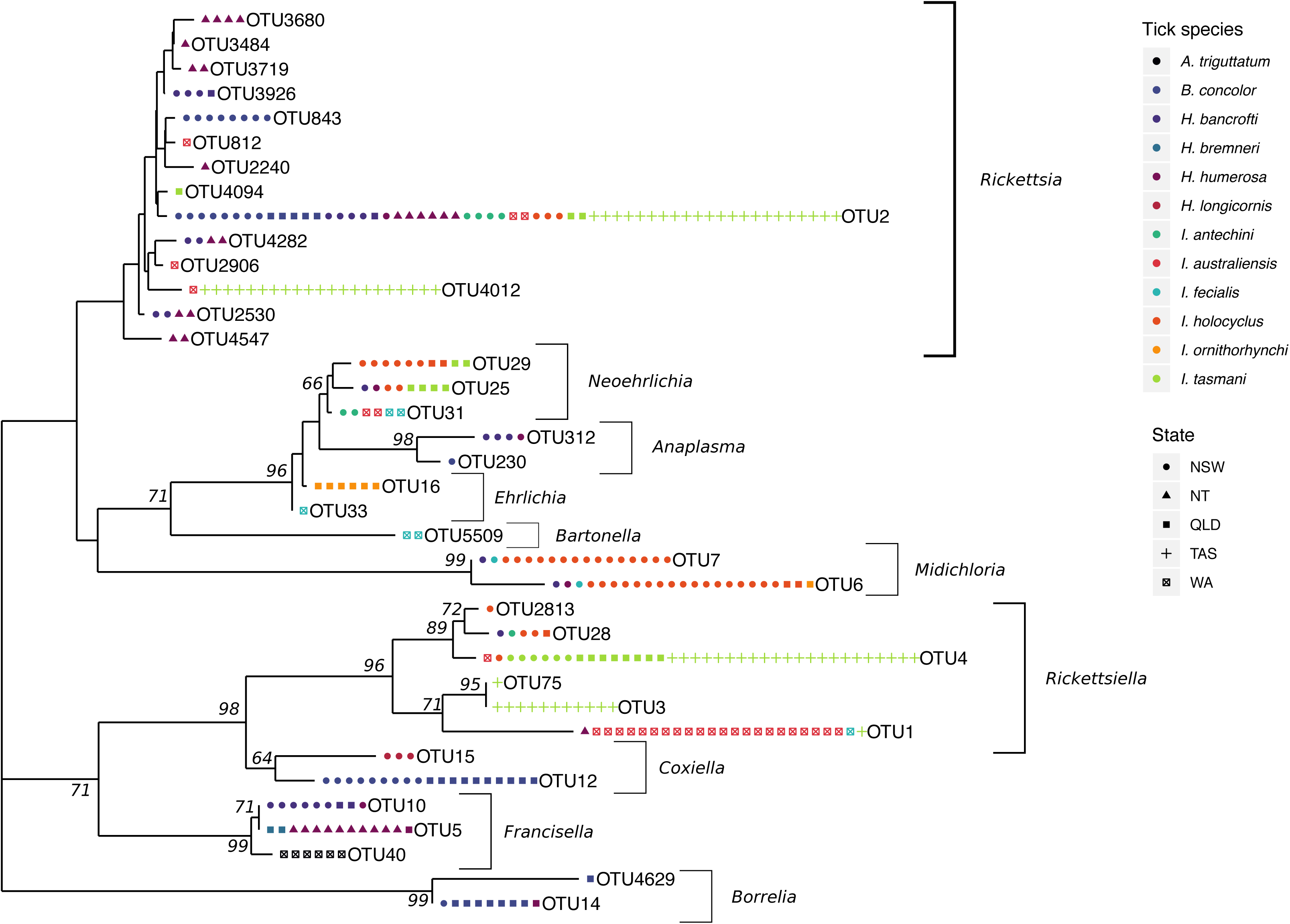
Neighbour-joining phylogenetic tree displaying taxa of interest prevalence and distribution among tick samples. Geographic data relating to tick collection is represented by state and territory; New South Wales (NSW), Northern Territory (NT), Queensland (QLD), Tasmania (TAS) and Western Australia (WA). Evolutionary histories were inferred based on the Tamura 3-parameter model with bootstrap analysis (10,000 replicates) (bootstrap values >60 are displayed). Tick samples were considered positive for taxa of interest if >100 sequences present. Information on number of sequences, taxonomic identity (as inferred from NCBI MegaBLAST analysis with nucleotide database), and sequences can be found in supplementary information (Supplementary File S4 & S5).

A proposed novel *Anaplasma* sp. (OTU230, MK814412, 96.3% identity) was identified in a single *B. concolor* (1/20) collected from (ex) echidna from NSW. A second *Anaplasma bovis*-like OTU (OTU312, JN862824, 97.3% identity) was identified in *Haemaphysalis bancrofti* (3/10) ex red-necked wallaby and long-nosed bandicoot and *Haemaphysalis humerosa* (n = 1/13) tick ex long-nosed bandicoot, all from NSW.

OTU5509 was assigned to the genus *Bartonella* and had a top BLAST hit of *Bartonella bacilliformis* (LN624026, 92.1% identity). OTU5509 was identified in two samples, however, was present in an extremely low number of sequences (2) in each case. Both samples were *I. fecialis* collected from two different quenda in WA. The low number of sequences means that in the case of statistical analysis, this OTU would have been filtered out. While there was no sufficient match to this sequence, it is noted that in many instances native Australian *Bartonella* species are lacking sequence information for this region of the 16S gene (V1-2).

Two OTUs (OTU14, CP025785, 100% identity; and OTU4629, CP025785, 100% identity) were identified as *‘Ca.* B. tachyglossi*’.* Sequences were identified in *Bothriocroton concolor* (8/20) ex echidnas from QLD & NSW and *H. humerosa* (1/13) ex northern brown bandicoot from QLD.

A *Coxiella*-like organism (OTU12, CP032542, 95.5% identity) was identified exclusively in *B. concolor* (19/20) ex echidna from NSW and QLD. This was the second most abundant OTU identified in *B. concolor*, accounting for ∼20% of the overall sequences. A *Coxiella* sp. (OTU15, KC170757, 100% identity) was identified in *Haemaphysalis longicornis* (3/4) ex red fox from NSW and, OTU represented ∼61.4% of the overall assigned bacterial sequences.

A novel *Ehrlichia* sp. (OTU33, AY309970, 96.3% identity) was identified in *I. fecialis* (1/13) ex quenda from WA. Other *Ehrlichia* OTUs (OTU16 and 1632) represented sequences from ‘*Ca.* Ehrlichia ornithorhynchi’ in *Ixodes ornithorhynchi* (6/10) ex platypus QLD and these sequences accounted for the majority of sequences (∼50.2%) from *I. ornithorhynchi*.

A *Francisella* endosymbiont (OTU5, AF001077, 98.0% identity) was identified in *H. humerosa* (11/13) ticks ex northern brown bandicoots from NT and QLD, and *Haemaphysalis bremneri* (2/4) ex possum (species unknown) in QLD. OTU5 was the most abundance sequence in *H. bremneri* and *H. humerosa*, representing 27.3% and 52.4% of the assigned sequences, respectively. A second *Francisella* endosymbiont (OTU10, AB001522, 99% identity) was identified in *H. bancrofti* (8/11) ex red-necked wallabies from QLD and NSW, and ex a long-nosed bandicoot in NSW; and in a single *H. humerosa* (1/13) ex red-necked wallaby from NSW. OTU10 was the most abundance sequences in *H. bancrofti* representing 27.0% of the assigned sequences. OTU40 (AF001077, 97.6% identity) was also identified as a *Francisella*-like endosymbiont from *A. triguttatum* (6/6) ex red kangaroo from WA. It was highly abundant in these ticks, accounting for 76.9% of the overall sequences.

*Midichloria* (OTU6, FM992372, 100% identity) was identified in *I. holocyclus* (20/36) ex long-nosed bandicoots from NSW and QLD, *I. fecialis* (1/13) ex red-necked wallaby NSW, *H. bancrofti* (1/11) ex long-nosed bandicoot NSW, and *H. humerosa* (1/13) ex long-nosed bandicoot NSW. *Midichloria* (OTU7, FM992373, 100% identity) was identified in *I. holocyclus* (15/36) ex long-nosed bandicoots from NSW and QLD, *I. fecialis* (1/13) ex red-necked wallaby NSW and *H. bancrofti* (1/11) ex long-nosed bandicoot NSW. Overall a total of 15 ticks had both OTU6 and OTU 7, which included *I. holocyclus* (13/36) ex long-nosed bandicoots from NSW and QLD, *I. fecialis* (1/13) ex red-necked wallaby NSW and *H. bancrofti* (1/11) ex long-nosed bandicoot NSW. While the CMm blocking primer was incorporated into the PCRs conducted on *I. holocyclus, Midichloria* sequence were still observed in 2/9 females, 3/6 males and 16/21 nymphs. *Midichloria* was the most abundant sequence from *I. holocyclus* accounting for 36.1% of assigned sequences.

Three *Neoehrlichia* OTUs were identified. ‘*Ca*. Neoehrlichia arcana’ (OTU25, KT203914, 99.3% identity) was identified in eight ticks, *I. tasmani* (4/51), *I. holocyclus* (2/36)*, H. bancrofti* (1/11) and *H. humerosa* (1/13), ex long-nosed bandicoots NSW and QLD. ‘*Ca*. Neoehrlichia australis’ (OTU29, KT803957, 100% identity) was identified in 10 ticks, *I. holocyclus* (8/36) and *I. tasmani* (2/51), ex long-nosed bandicoots NSW and QLD. Three ticks (*I. tasmani* and *I. holocyclu*s) were co-infected with both ‘*Ca*. N arcana’ and ‘*Ca*. N. australis’. A novel *Neoehrlichia* species (OTU31, MG670107, 97.3% identity) was identified in *I. fecialis* (2/13) and *I. australiensis* (2/26) ex quenda from WA, and *I. antechini* (2/29) ex brown antechinus from NSW.

Due to the conserved nature of the *Rickettsia* genus at the 16S gene, reported previously (Roux and Raoult, 1995; Stothard and Fuerst, 1995), resolution to species level was not possible using the sequences obtained in the present study. *Rickettsia* sp. (OTU2, KF318168, 99.7% identity) had a widespread distribution among samples and was detected in 61 ticks *(B. concolor, H. bancrofti, H. humerosa, I. antechini, I. australiensis, I. holocyclus* and *I. tasmani*) from three states and one territory (NSW, NT, QLD, WA). *Rickettsia* was the most abundant sequence from *B. concolor* and *I. antechini* ticks accounting for 63.3% and 12.0% of the assigned sequences, respectively A *Rickettsiella*-like sp. (OTU1, LC388767, 92.7% identity) was identified in *I. australiensis* (22/26) and was the most abundant sequence accounting for 76.8% of assigned sequences. A *Rickettsiella*-like sp. (OTU3, EU430251, 95.1% identity) was identified in *I. tasmani* (11/51) ex Tasmanian devils and eastern barred bandicoot from TAS. A *Rickettsiella*-lik*e* sp. (OTU4, U97547, 98.4% identity) was identified in, *I. tasmani* (36/51) ex bandicoots, possum and a sugar glider from NSW, TAS and QLD; *I. australiensis* (1/26) ex western grey kangaroo from WA; and *I. holocyclus* (1/36) ex long nosed bandicoot NSW. OTU4 accounted for 30.2% of the overall sequences assigned for *I. tasmani*.

## 4. Discussion

Currently there is a scarcity of detailed information about the life cycles of Australian ticks, however it is generally assumed that, as with other hard tick species around the world, the majority will exhibit a three-host life cycle (Barker and Walker, 2014). Despite these limitations, the opportunistic sampling strategy used in this study provides an economical way to survey a wide range of tick and wildlife fauna across a range of geographical areas across Australia. The paucity of larvae from this data set is also a limitation of this type of sampling. Tick larval stages, which may indicate the presence of transovarially-transmitted organisms (Kwan et al., 2017), are difficult to see with the naked eye, and in situations where collection of ticks is not the main objective, they are easily overlooked (Lydecker et al., 2019). Furthermore, the opportunistic nature of the sampling precluded assessment of the infestation rates of the various hosts. In future, if TBPs of humans and the wildlife themselves are identified, further extensive surveys of tick-pathogen-wildlife ecologies will require more targeted and comprehensive approaches in order to gather sufficient relevant epidemiological data.

As with previous studies (Swei and Kwan, 2016; Zolnik et al., 2016) alpha diversity measures were highly variable both within and between tick species. This may be driven by a variety of factors such as; starting material (i.e. size/life stage of tick), extraction methods, library preparation, normalisation of DNA input concentrations for sequencing, batch sequencing effects and bioinformatics (Greay et al. 2018). Within the present study, sequencing depth was likely most impacted by sample input type and normalisation of DNA concentrations, as all samples went through the same library preparation and bioinformatic analysis. Overall, bacterial diversity of ticks started to plateau by 50,000 sequences however, it is noted that in some cases this plateau was not achieved and deeper sequencing would be required in order to confidently characterise the full suite of bacterial taxa present. This was most evident in *A. triguttatum*, *B. auruginans*, *I. trichosuri* and *I. ornithorhynchi* and therefore we recommend a minimum of 100,000 sequences per sample for future bacterial 16S amplicon studies in these species to be confident that the complete bacterial community has been sampled. Studies that investigate the shift of the core microbiome (i.e. most abundant bacteria taxa) relative to a given parameter may not require the same depth of sequencing, as seen in previous studies investigating the effect of temperature (Thapa et al., 2019) and life stage (Andreotti et al., 2011).

After bioinformatic analysis, including stringent quality filtering, a large number of OTUs remained with relatively low number of reads. This has been noted in previous studies of the tick microbiome (Budachetri et al., 2016; Zolnik et al., 2016) and in the broader field of high-throughput microbiome studies (Pollock et al., 2018). Depending on bioinformatic analysis and quality filtering, the number of OTUs and therefore diversity can vary greatly among studies (Greay et al., 2018). In addition, current practice in microbiome studies is to normalise count data, however models show that this can oversimplify the data (McMurdie and Holmes, 2014).

Ordination analysis demonstrated that tick species was the strongest predictor of bacterial composition. Beta-diversity analysis showed that the common marsupial tick, *I. tasmani*, exhibited a variable bacterial composition. This diversity may be explained by the wide geographic distribution of this tick in Australia, its own high genetic diversity (Burnard and Shao, 2019), and its ability to parasitise many marsupial species. In the present study, bacterial profiling of *I. tasmani* included ticks from three states (NSW, QLD, TAS) and seven host species. With this in mind, we suggest that future studies should combine careful taxonomic status identity with data on microbial communities of *I. tasmani*. Although sample sizes were limited in ‘host-specialist’ ticks *B. auruginans* (wombats)*, H. bremneri* (possums)*, I. antechini* (antechinus) and *I. ornithorhynchi* (platypus), these specimens showed less diversity of bacterial communities between samples.

An important caveat on the relative diversity of bacterial communities in the present study was the use of a blocking primer inhibiting the amplification of ‘*Ca*. M. mitochondrii’. The inclusion of this blocking primer it vital to explore the full bacterial community of *I. holocyclus* (see Gofton et al., 2015), however it does impact the analysis and interpretation of the data. Due to the inhibition of this bacteria, there is an inherent bias in the bacterial community, and comparisons in alpha and beta-diversity measures must account for this. In the case of highly abundant organisms, the use of a blocking primer assay will likely not completely inhibit amplification, and thus a reduced level of the organisms may be still detected, as was the case in the present study. In this instance, the use of diversity measures that rely on presence/absence data will not be affected by this granted there is still some level of detection (i.e. number of species observed for alpha diversity and Jaccard index for beta-diversity). The use of more advanced, and usually preferred, diversity indexes may be impacted by the manipulation of bacterial composition, and care must be taken when comparing results with other studies (Greay et al., 2018). Despite these limitations, the use of blocking primers has shown to be vital in the context of investigating known tick-borne pathogens and novel related taxa from tick samples. In the present study we have characterised the bacterial communities of ten tick species for the first time. This information provides the first step of focusing future research and where this established technique should be applied.

Members of the Proteobacteria represented the most diverse and abundant (as determine by number of sequences) in Australian hard ticks analysed in the present study. This finding is consistent with previous microbiome studies from tick species, such as *Ixodes scapularis* (Sperling et al., 2017; Thapa et al., 2019), *Ixodes persulcatus* (Zhang et al., 2014; Kurilshikov et al., 2015) and *Amblyomma americanum* (Fryxell and DeBruyn, 2016). Additionally the identification of novel bacterial taxa in native Australian ticks species is consistent with recent findings (Gofton et al., 2015b; Panetta et al., 2017).

Currently in Australia, the ecology, epidemiology, and incidence of human TBDs remains largely a matter of conjecture, having received little scientific study compared with many other parts of the world (Graves and Stenos, 2017). Despite significant national interest, including a federal government senate inquiry (Radcliffe et al., 2016), the prevailing scientific opinion concludes that Lyme borreliosis (caused by *B. burgdorferi* sensu lato), for example, is absent from Australia (Chalada et al., 2016; Irwin et al., 2017). In the unique Australian environment, long isolated in geological terms, it is likely that unidentified tick-host life cycles have evolved, given the endemic wildlife (including tick fauna) present (Long, 2017; Beati and Klompen, 2019). It is possible that these cycles may contribute to zoonotic illness when humans encroach these sylvatic ecologies and become exposed to native ticks. Indeed, it is well documented that in endemic areas in Europe and North America, the causative agent of Lyme borreliosis is readily identified in wildlife (such as white-tailed deer, *Odocoileus virginianus*, and white-footed mice, *Peromyscus leucopus*) as well as their natural tick species (Bosler et al., 1984). The present study therefore is one of the first to explore the concept that Australian wildlife ticks should be a promising source to identify exotic and novel TBPs, and we focussed our search towards taxa known to be associated with TBDs overseas (‘taxa of interest’), namely *Anaplasma*, *Bartonella*, *Borrelia*, *Coxiella*, *Ehrlichia*, *Francisella*, *Midichloria*, *Neoehrlichia*, *Rickettsia* and *Rickettsiella*.

A new genetic variant of *Anaplasma bovis* was recently described from questing *A. triguttatum* ticks in WA and the nearby Barrow Island (Gofton et al., 2017). In the present study we report a range expansion of a genetically similar *A. bovis* along the east coast of Australia from two widespread *Haemaphysalis* species. As suggested by Gofton et al., (2017), this finding further supports the hypothesis that it is likely that endemic *A. bovis* genotypes exist in sylvatic cycles within native Australian ticks and wildlife fauna. Further research on the phylogenetic position of these *A. bovis* sequences is needed to understand their likely evolutionary history and relatedness to *A. bovis* genotype Y11 identified in Western Australia. While three other species of *Anaplasma* (*A. marginale*, *A. centrale* and *A. platys*) have been introduced to Australia (Rogers and Shiels, 1979; Callow, 1984; Angus, 1996), the present study did not identify any of these species in ticks from wildlife. This is likely due to the absence of the cattle tick (*R*. (*B*.) *australis*) and the brown dog tick (*R. sanguineus*) specimens from wildlife hosts in the present study. A species of *Anaplasma* identified exclusively in *B. concolor* from echidnas sheds more light on the diverse range of microbes that have been described from this specialist tick (Loh, 2018).

Although the presence of a *Bartonella* sp. was identified by only four sequences, its absence in control samples means that this likely represents a true finding in female *I. fecialis*, which were parasitising different individuals of the same host (quenda) in south-west WA. Recent studies have shown that Australian marsupials and native rodents harbour a range of distinct *Bartonella* species; ‘*Ca*. Bartonella antechini’ has been identified in ticks (*I. antechini*) and fleas (*Acanthopsylla jordani*) from mardo (*Antechinus flavipes*) in south-west WA (Kaewmongkol et al., 2011c), ‘*Ca*. Bartonella woyliei’ in woylie ticks (*I. australiensis*) and fleas (*Pygiopsylla hilli*) in south-west WA, and ‘*Ca*. Bartonella bandicootii’ in fleas (*Pygiopsylla tunneyi*) from western barred bandicoots (*Perameles bougainville*) on Bernier and Dorre Island (Kaewmongkol et al., 2011a). Reports of *Bartonella* spp. occurring outside south-west WA include *Bartonella australis* ex eastern grey kangaroos (*Macropus giganteus)* (Fournier et al., 2007). Molecular detection of *Bartonella* DNA from Australian ticks has also been reported in ticks (*I. tasmani*) parasitising koalas (*Phascolarctos cinereus*) from Philip Island, VIC (Vilcins et al., 2009b). Additional studies on *Bartonella* in Australia wildlife include those by Gundi et al. (2009), Kaewmongkol et al. (2011a) and Dybing et al. (2016). Due to the limited size and region of the 16S amplicon generated in the present study, there were insufficient relevant reference sequences available from other Australian *Bartonella* species for comparison. A study by Kaewmongkol et al. (2011b) into flea-derived *Bartonella* from native and introduced Australian species suggests co-evolution of marsupial hosts, their fleas and the *Bartonella* species.

‘*Ca*. Borrelia tachyglossi’ was identified in 8/20 *B. concolor* ticks from echidnas in QLD and NSW, an anticipated finding given previous research into this organism (Loh et al., 2016). Importantly however, the present study provides the first evidence of ‘*Ca*. B. tachyglossi’ sequences from a female *H. humerosa* tick parasitising a northern brown bandicoot from QLD. In addition, no *Bothriocroton* ticks were present in the extraction batch of the positive *H. humerosa* sample, and there was no evidence of *Borrelia* sequences from any controls. Despite the wide geographical range of *H. humerosa* ticks, the restricted finding of ‘*Ca*. B. tachyglossi’ from QLD supports Loh et al. (2016), suggesting a restricted geographical distribution along the east coast of Australia.

In tick microbiome studies overseas, *Coxiella* spp. are commonly identified (Khoo et al., 2016; Machado-Ferreira et al., 2016). Additionally, recent studies on Australian ticks have also identified *Coxiella* spp. in *A. triguttatum* (Cooper et al., 2013; Gofton et al., 2015b), *B. auruginans* (Vilcins et al., 2009c), *I. holocyclus* (Cooper et al., 2013) and *R. sanguineus* (Oskam et al., 2017). Interestingly the present study did not identify the widespread presence of *Coxiella* spp. with only two unique OTUs identified from *B. concolor* and *H. longicornis*, both of which appear to be host specific to those tick species, and the causative agent of Q-Fever (*C. burnetii*) was not identified from ticks in the present study.

*Ehrlichia* sequences (‘*Ca*. E. ornithorhynchi’) from platypus ticks, *I. ornithorhynchi*, were recently described (Gofton et al., 2018). In the present study, ‘*Ca*. E. ornithorhynchi’ was exclusively observed within *I. ornithorhynchi,* suggesting this genetically distinct *Ehrlichia* species has a unique and host-specific relationship with the platypus. During this study, a potentially novel species of *Ehrlichia* was identified in an *I. fecialis* female tick from a quenda in WA. BLAST results show it is a close relative to an *Ehrlichia* sp. detected in *Haemaphysalis* ticks from Japan (Inokuma et al., 2004). No sequences from the recently described ‘*Ca*. E. occidentalis’ (Gofton et al., 2017) were identified in the present study, however it is noted *A. triguttatum* were only represented by six samples.

*Francisella*-like endosymbionts have been widely reported in ticks overseas, such as *Dermacentor* spp. (Scoles, 2005), *Dermacentor occidentalis* (Gurfield et al., 2017), *Haemaphysalis longicornis* (Wang et al., 2018) and *Hyalomma rufipes* (Szigeti et al., 2014), and *Francisella* sequences have previously been identified in *A. fimbriatum* ticks from reptiles in the Northern Territory of Australia (Vilcins et al., 2009a). In the present study, a *Francisella*-like endosymbiont was identified in 100% of *A. triguttatum* ticks, and in a high proportion. The high prevalence and relative abundance of the *Francisella*-like organisms in *A. triguttatum* may be of medical relevance with respect to this common human-biting tick, particularly given the recent report of *Francisella* bacteraemia in WA (Aravena-Román et al., 2015). Importantly, as noted previously for *I. holocyclus* ticks, organisms in lower abundance may be masked by abundant endosymbionts, unless samples are sequenced deeply and/or a blocking primer is used (Gofton et al., 2015a). Therefore, future bacterial profiling studies of *A. triguttatum* should take these factors into consideration.

While ‘*Ca*. M. mitochondrii’ has been documented in *I. holocyclus,* the present study provides the first report of ‘*Ca*. M. mitochondrii’ within additional species of native Australian ticks (*I. fecialis, H. bancrofti* and *H. humerosa*). Despite this expansion among tick species, identification of ‘*Ca*. M. mitochondrii’ remains confined to NSW and QLD as previously reported (Gofton et al., 2015a). ‘*Ca*. M. mitochondrii’ has been detected overseas in laboratory models (Cafiso et al., 2019), wildlife (Serra et al., 2018), and humans (Mariconti et al., 2012). Despite the use of a CMm blocking primer by Gofton et al. (2015a) in the library preparation of *I. holocyclus*, the bacteria was still identified in the majority of ticks (21/36); however its abundance was greatly reduced to an average of 26.1% of reads per sample in comparison to previous research demonstrating a relative abundance of 98.2% (Gofton et al., 2015a).

A recent addition to the Anaplasmataceae family, ‘*Ca*. Neoehrlichia mikurensis’ was first isolated from wild rats and their ticks (*Ixodes ovatus*) in Japan (Kawahara et al., 2004). In Europe, the hedgehog (*Erinaceus* spp.), a common peri-urban dweller, has been shown to play an important role in the life cycle of ‘*Ca*. N. mikurensis’ (Földvári et al., 2014; Jahfari et al., 2017). A high proportion of bandicoot ticks were infected with *Neoehrlichia* spp. in this study, suggesting that these marsupials may play an important role in the life cycle of this bacterium on the Australian continent. Bandicoots also frequently inhabit gardens and have relatively close contact with humans (Carthey and Banks, 2012). Closer study of these marsupials may provide vital information about the life cycle of these microbes and the potential risk for human infection.

*Rickettsia* spp. are among the only currently recognised TBPs affecting people in Australia, however, as previously outlined, the conserved nature of the 16S gene in *Rickettsia* precludes rigorous species delimitation from the relatively short sequences generated in this dataset. In addition to the known human pathogens (*R. australis* and *R. honei*) (Graves and Stenos, 2017), the presence of *Rickettsia* has been demonstrated from a variety of Australian tick species. *Rickettsia gravesii* has been described from *A. triguttatum* (Li et al., 2010; Abdad et al., 2017) and *Rickettsia* sequences have been previously identified in *I. tasmani* ex Tasmanian devils (Vilcins et al., 2009c) and *Amblyomma fimbriatum* from reptiles (Vilcins et al., 2009a).

The diversity and significance of the *Rickettsiella* genus remains largely unknown globally. Phylogenetic analysis of sequence data places the genus within the Coxiellaceae family (Fournier and Raoult, 2005; Leclerque and Kleespies, 2012). While the presence of these organisms has been well documented in tick species around the world and in some cases, they have been identified as pathogenic to arthropods (Kurtti et al., 2002; Leclerque et al., 2012), disease causation within the vertebrate hosts remains unknown at the present time. A study by Vilcins et al. (2009c) identified *Rickettsiella* in *I. tasmani* ex koalas from Phillip Island, just off the coast of south eastern Australia. A genetically similar sequence was identified in the present study (OTU3) from *I. tasmani* exclusively in Tasmania.

While this is the first study to characterise the bacterial communities within native Australian wildlife ticks, it is apparent that much research is still required in order to better understand the tick-associated microbial life cycles and their ecologies, let alone elucidating their potential for transmission and pathogenicity in vertebrate hosts, including humans. Furthermore, whilst cost-effective, ticks were collected opportunistically resulting in an inherent geographical bias, confining sampling largely to urban areas along the east coast of the Australian continent. Nevertheless, this area is also where most humans receive tick bites, although reliable data about this is also lacking. In addition, the present study suggests that ‘taxa of interest’ are largely restricted to a combination of geographical location and tick species. For example, ‘*Ca*. Midichloria’ was identified in *H. humerosa* and *I. fecialis* from NSW only and was absent in samples from NT for *H. humerosa* and WA and TAS for *I. fecialis*.

The analysis of ticks removed from wildlife hosts comes with the inherent complication of a blood meal. The host blood meal has been recently shown to influence both the tick microbiome composition and the presence of pathogens (Landesman et al., 2019; Swei and Kwan, 2017). Further targeted studies are needed to assess the source of microbes with respect to a tick or host origin (Irwin et al., 2018). In addition, limitations in the current sequencing technologies mean that high-throughput methods favour short amplicons which may not be able to accurately discriminate bacterial species, as is the case with *Rickettsia* in particular (Gihring et al., 2012). Furthermore, important caveats to consider in metabarcoding microbial diversity analysis include PCR efficiency (including primers used and length of target amplicon), variations in 16S gene copy number (Ahn et al., 2012), sequencing depth, machine cross-talk, bioinformatic analysis and taxonomic assignment. While there have been some recent attempts to correct these biases (Kembel et al., 2012; Rosselli et al., 2016), their application to uncharacterised metagenomic samples remains limited. While 16S metabarcoding continues to be vital in bacterial biodiversity discovery (Carpi et al., 2011; Tessler et al., 2017), future molecular studies incorporating metagenomics and metatranscriptomics, to detect actively expressed genes (Cabezas-Cruz et al., 2018; Greay et al., 2018), will be useful in characterising the full suite of micro-organisms present in Australian ticks. A multi-disciplinary approach incorporating cell culture, in vitro tick studies and morphological techniques (e.g. fluorescence microscopy) will also be needed to assess the potential for transmissibility and pathogenicity of these novel tick-associated organisms.

## 5. Conclusions

With over 75% of emerging human infectious pathogens originating from wildlife (King, 2014), surveillance methods that target these species are important for investigations of emerging and exotic infectious disease. Research into the interactions of wildlife hosts, ticks and pathogens in Europe and North America continues to highlight the complexity of these dynamic systems (Ostfeld et al., 2018; Tomassone et al., 2018). Results from the present study build on recent research into Australian tick-associated microbes, further highlighting the diversity of organisms present that appear related, yet distinct from their overseas counterparts. With the evolutionary history of Australia’s unique tick species (Beati and Klompen, 2019) and wildlife fauna, it is likely that reports of their tick-associated microbes (potential pathogens) will continue to reveal taxonomic differences from those described in the northern hemisphere. As such, future research on emerging tick-borne zoonoses should include methods that are able to detect novel micro-organisms in humans and reservoirs and include a variety of sample types such as blood, tissue (e.g. skin, spleen etc.) and tick from both the host and environment (questing). As anthropogenic changes to the environment continue to grow in Australia, a greater emphasis on wildlife disease surveillance is critical to ensure the early detection of potential infectious diseases affecting humans, livestock, companion animals and wildlife (Woods et al., 2019). Advances in pathogen detection and characterisation are greatly enhanced by collaboration; the authors advocate for continued multidisciplinary efforts between health professionals, researchers, land managers and local communities.

## Supporting Information

### Data availability

Associated metadata output from bioinformatic analysis and subsequent data visualisation is available on FigShare repository https://doi.org/10.6084/m9.figshare.c.4608803.v1. Next-generation sequencing data can be accessed from NCBI Sequence Read Archive under BioProject: PRJNA559059 (BioSample accession numbers: SAMN12512474 –SAMN12512711).

## Acknowledgements

The authors would like to thank collaborating research teams, veterinary clinics, wildlife centres and members of the public for contributing samples to this study. Supplementary File S1 contains complete list of collectors and institutions. This study was part-funded by the Australian Research Council (LP160100200) and Bayer HealthCare (Germany) and Bayer Australia. S.E. received funding from a Murdoch University honours scholarship. S-M.L. was support by a Murdoch International Postgraduate scholarship. The authors also acknowledge the Pawsey Supercomputing Centre and the Western Australia State Agriculture Biotechnology Centre for use of its facilities. We thank Dr. Alexander Gofton and Ms. Telleasha Greay for advice on next-generation sequencing methods. The funders had no role in the experimental design, data collection and analyses.

## Compliance with ethical standards

This study was conducted under the compliance of the Australian Code for the Responsibility Conduct of Research (2007) and Australian Code for the Care and Use of Animals for Scientific Purposes, 2013. Tick collection was carried out opportunistically with the approval from the Murdoch University Animal Ethics Committee.

## Supplementary Files

**Supplementary File S1**. Sample metadata of tick-wildlife records identified from the present study.

**Supplementary File S2.** Sequence information from 16S rRNA bacterial profiling of ticks on the Illumina MiSeq platform after bioinformatic analysis; (a) number sequences assigned to 97% operational taxonomic units (OTUs) and (b) number of sequences per sample.

**Supplementary File S3.** Principal coordinate analysis (PCoA) plot of the weighted UniFrac distance matrix. Plots show beta diversity of tick bacterial communities, displayed by tick species and life stage. Operational taxonomic units (OTUs) that represented <100 sequences in a tick sample were removed.

**Supplementary File S4.** Number of sequences relating to the taxa of interest in tick samples. The top hit from NCBI BLAST results showing GenBank accession number, percent identity and E-value are given, and final evaluation of taxonomic assignment.

**Supplementary File S5.** Fasta sequences for operational taxonomic units (OTUs) used in the taxa of interest investigation.

## Notes

#### Summary of Updates

Additions to methodology and discussions. Two figures moved to supplementary information and one figure removed from the manuscript.

https://figshare.com/collections/Wildlife_tick_microbiome/4608803/1

