## Supplementary figures and images for "Bacterial community profiling highlights complex diversity and novel organisms in wildlife ticks"

### Supplementary File S2

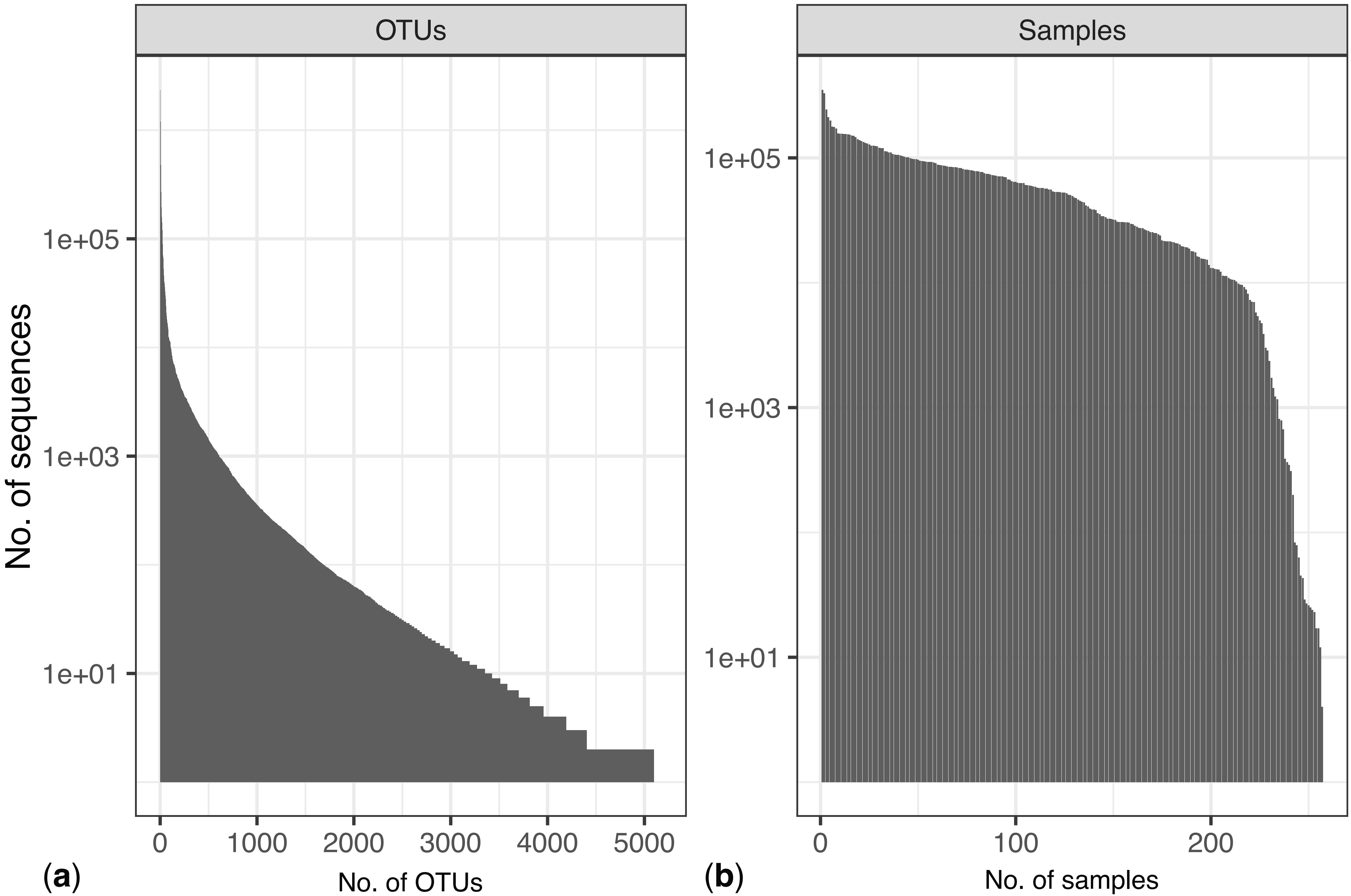

### Supplementary File S3

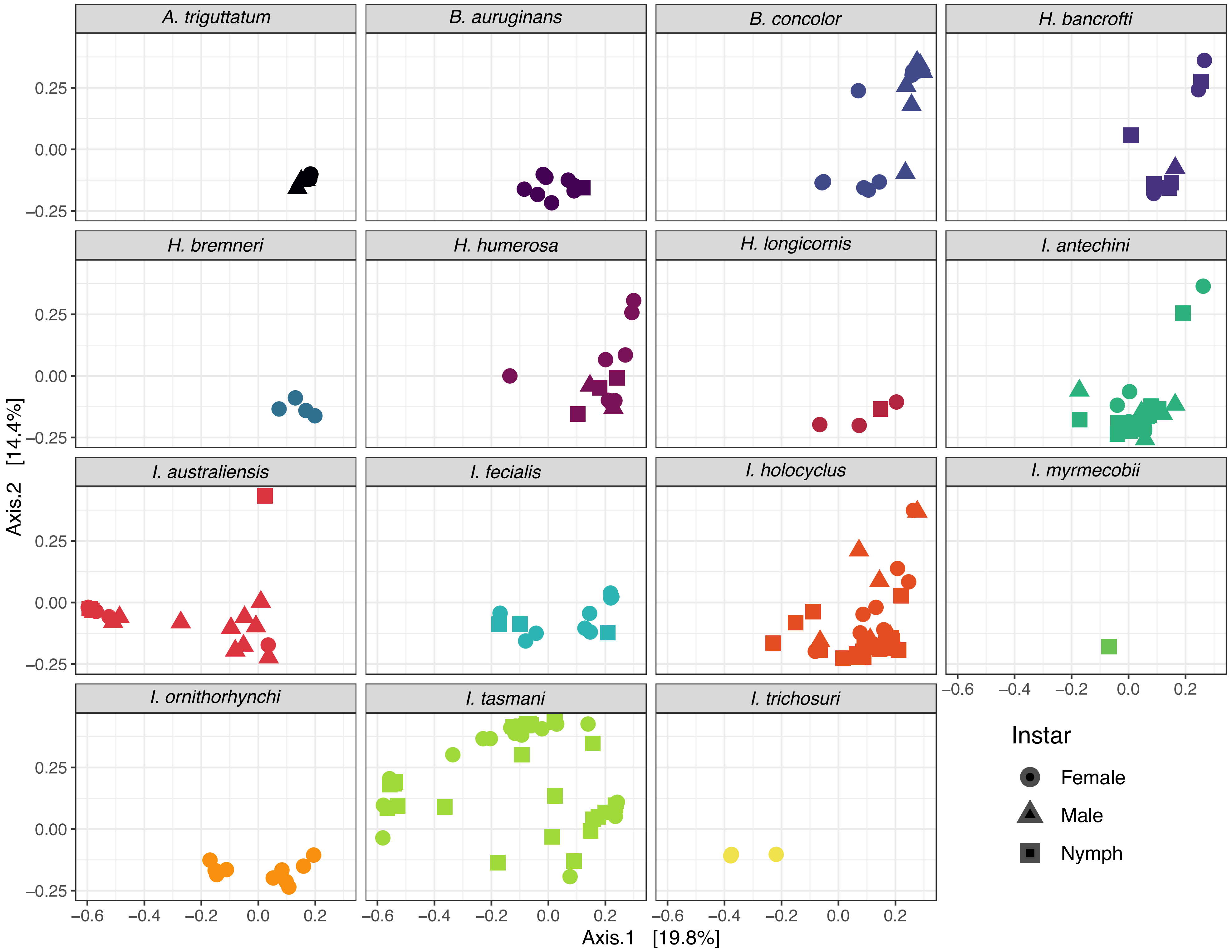
